# Inferring future changes in gene flow under climate change in riverscapes

**DOI:** 10.1101/2022.05.10.491062

**Authors:** Souta Nakajima, Hiroaki Suzuki, Makoto Nakatsugawa, Ayumi Matsuo, Shun K. Hirota, Yoshihisa Suyama, Futoshi Nakamura

**Affiliations:** Graduate School of Agriculture, Hokkaido University, Kita-ku N9W9, Sapporo, Hokkaido 060-8589, Japan; Graduate School of Engineering, Muroran Institute of Technology, Mizumoto-cho 27-1, Muroran, Hokkaido 050-8585, Japan; Research Institute of Energy, Environment and Geology, Hokkaido Research Organization, Kita-ku N19W12, Sapporo, Hokkaido 060-0819, Japan; Graduate School of Agricultural Science, Tohoku University, Yomogida 232-3, Naruko-onsen, Osaki, Miyagi 989-6711, Japan

**Keywords:** model-based riverscape genetics, cold-water fish, *Cottus*, water temperature, global warming

## Abstract

Global climate change poses a significant threat to the habitat connectivity of cold-water-adapted organisms, leading to species extinctions. Because gene flow is itself a functional connectivity among wild populations, if gene flow is modeled by landscape variables, changes in population connectivity could be predicted. In this study, using a model-based riverscape genetics technique and a hydrological model to estimate water temperature, we inferred the determinants of and future changes in gene flow of the fluvial sculpin *Cottus nozawae* in the upstream section of the Sorachi River, Hokkaido, Japan. As a result of model selection, stream order, water temperature, slope, and distance were detected as landscape variables affecting the strength of gene flow in each stream section. In particular, the trend of greater gene flow in sections with higher stream order and lower temperature fluctuations or summer water temperatures was pronounced. The map from the model showed that gene flow is overall prevented in small tributaries in the southern area, where spring-fed environments are less prevalent. Estimating future changes in gene flow using future water temperature predictions, genetic connectivity was predicted to decrease dramatically until the end of the 21st century under IPCC representative concentration pathway scenario 8.5 (RCP8.5).

## Introduction

Global climate change modifies water temperatures and flow regimes, the two key habitat factors affecting freshwater species, posing a critical threat to stream ecosystems [1]. The spatial distribution of species’ suitable habitats shifts with environmental changes, and population fragmentation due to impassable environments may eventually result in local and/or species extinctions [2]. Numerous studies have predicted changes in species distributions and suitable habitats of stream organisms [3], but how will the actual population connectivity and migration potential change?

Gene flow represents the valid connectivity of wild populations and has an important function in species viability [4]. The strength of gene flow is usually discussed individually from the observed genetic structure, but if gene flow could be modeled by landscape variables, the gained knowledge regarding gene flow could be generalized and used to predict its future changes. The relationships between gene flow and landscape variables have been investigated in the field of landscape genetics [5]. However, most analytical techniques developed in landscape genetics exert only poor power in linear and dendritic stream ecosystems [6,7], making it difficult to predict future changes in gene flow in riverscapes. Even in streams, regression models can be created by contrasting a genetic distance matrix against pairwise differences in local conditions [8], but this approach fails to account for the network architecture and for the attributes in all the spaces that individuals must pass through when traveling between sampling sites [7,9]. Another versatile approach to investigating the effects of landscape elements on gene flow is defining “landscape resistance” surfaces and assessing the relationship between genetic distance and cumulative resistance between populations (isolation by resistance; IBR [10]). Although this idea has been applied to studies on stream ecosystems in a couple of cases [11,12], the landscape resistance must be parametrized *a priori* through expert opinion or other empirical methods (e.g., using the inverse of species distribution model estimates) [13]. To understand the real gene flow, its determinants should be identified directly from genetic data. Fortunately, alternative methods for modeling gene flow from genetic data within a spatially explicit graph-theoretic framework have been developed rapidly in recent years [6,9]. Although not yet practically applied to predictions under environmental changes, we thought that these “riverscape genetics”-dedicated methods are the key to determining landscape resistance and modeling current and future gene flow.

Another theme that makes riverscape genetics challenging is the data availability of key environmental elements such as water temperatures. For terrestrial organisms, globally available climate data such as WorldClim [14] are commonly used to estimate the effects of climate change. However, data on current and future water temperatures that are critical for stream organisms are difficult to obtain as well as data on temporal flow rate. Although some studies have used air temperature data as a surrogate of water temperatures [15], water temperatures do not actually coincide with air temperatures. In particular, local spatial heterogeneity in water temperatures caused by groundwater discharge and other factors is truly a source of ecosystem diversity and resilience to climate change that cannot be ignored [16–19]. Therefore, it is critical in riverscapes to utilize water temperature information considering the spatial heterogeneity generated by hydrogeological factors.

*Cottus nozawae* is a cold-water-adapted sculpin inhabiting northern Japan. Since the distribution and ecology of this species are highly influenced by summer water temperatures [20], available habitats are expected to decrease significantly under climate change [16]. At a local scale, streams with low summer water temperatures characterized by spring-fed environments have been shown to display high population densities and to be the source of individuals in a watershed [17,21]. Under ongoing climate change, the migration of this species is expected to be frequently blocked by unsuitable habitats, resulting in population fragmentation and shrinkage. Understanding population connectivity in cold-water fishes is important for conservation and is a hot topic in riverscape genetics [22].

Considering the challenges of data availability and analysis in riverscape genetics, we thought that the recently developed model-based riverscape genetics approaches and process-based hydrological model to estimate water temperatures will enable the modeling and future prediction of gene flow in cold-water fish. The aims of this study are (i) to identify the factors determining the gene flow of *C. nozawae* in the stream network, (ii) to model the strength of gene flow using landscape variables and predict its future changes, and (iii) to discuss the applicability of riverscape genetics modeling in conservation ecology.

## Material and Methods

In 2019, 376 individuals of *C. nozawae* were captured by electrofishing from 13 sites located in the upstream section of the Sorachi River, Hokkaido, Japan (figure S1; table S1). Because no river-crossing structures that would obviously prevent fish migration are present between sampling sites, this area is considered suitable for evaluating the effects of landscape variables. Regarding the environmental conditions, the tributaries in the northern volcanic watersheds have spring-fed environments with stable water temperatures and flow regimes [16,17]. The stream network was viewed as a graph consisting of 24 “nodes” and 23 “edges” (figure 1a). We defined “nodes” as the sampling sites and major tributary confluences between them, and gene flow modeling was conducted with “edges”, the stream segments between adjacent nodes, as units.

**Figure 1.**
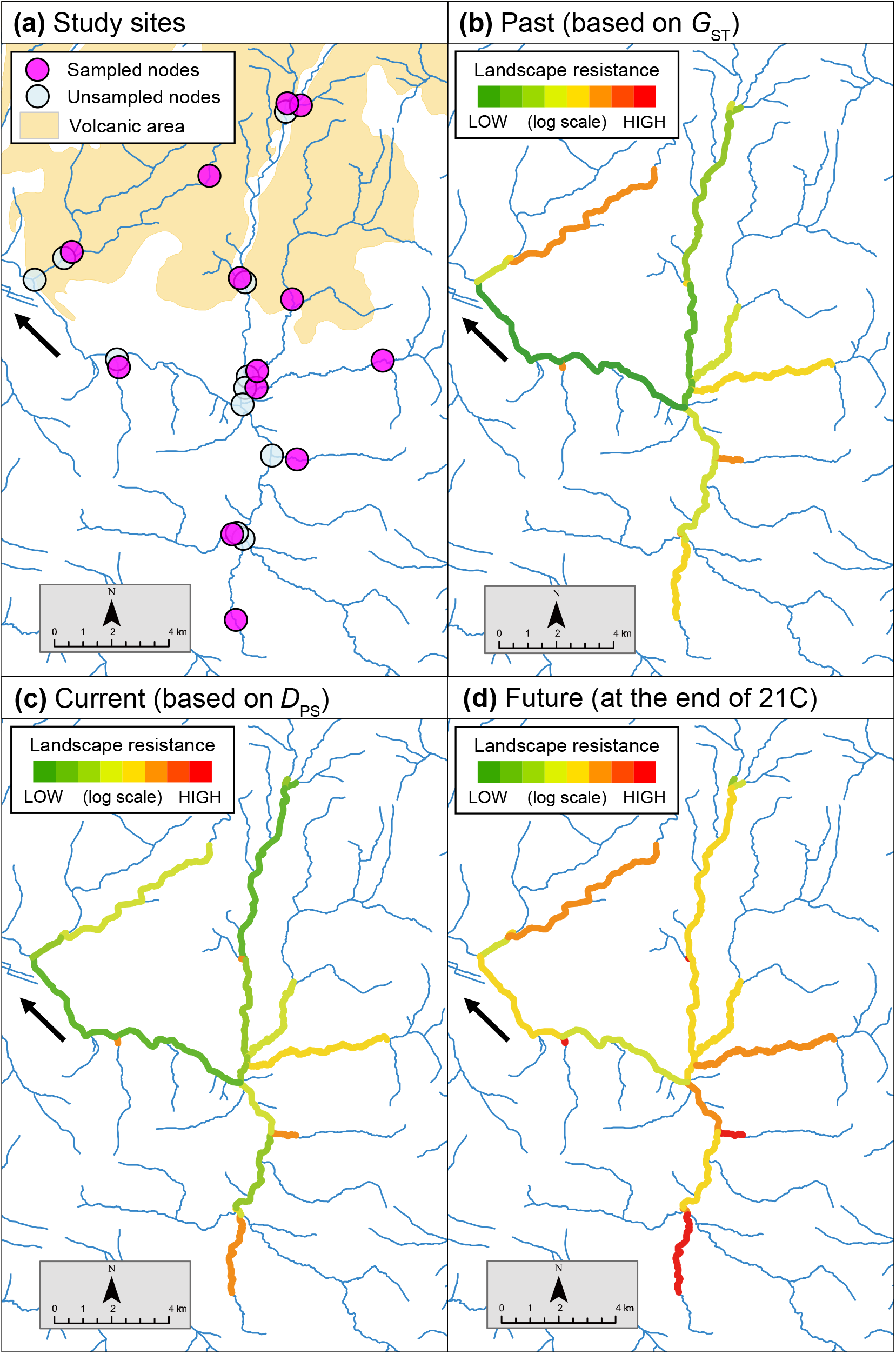
Maps of the Sorachi River watershed showing the (a) study sites represented as a graph and (b–d) landscape resistance estimated by BGR models. Three patterns are shown: (b) long-term gene flow modeled by *G*_ST_, (c) recent gene flow modeled by *D*_PS_, (d) predicted gene flow at the end of the 21st century derived by substituting future water temperatures into the model derived by *D*_PS_.

From the DNA extracted from the fin tissue of each individual, 212 SNPs were gained from the multiplexed ISSR genotyping by sequencing (MIG-seq) method [23,24] and subsequent data filtering by STACKS 2.41 [25] (see appendix 2 for details). We probabilistically modeled the relative migration rate (edge passability) of each edge as a function of 10 possible landscape variables (table 1, figure S2) using the bidirectional geneflow in riverscapes (BGR) model [9]. This is a novel method that can model bidirectional gene flow in stream networks using genetic distance matrices as responses and landscape variables as explanatory variables, rigorously accounting for the spatial autocorrelation structure of stream networks. As genetic distances, we used *G*_ST_ [26] as an indicator of historical gene flow, and *D*_PS_ (1-the proportion of shared alleles [27]) as an indicator of current gene flow [28,29]. Landscape variables related to flow rate and water temperature were estimated by a hydrological model based on Suzuki H et al. [30], which considers differences in groundwater discharge depending on catchment geology (see appendix 3 for details). The other variables were quantified from the National Land Numerical Information from the Ministry of Land, Infrastructure, Transport and Tourism (MLIT) of Japan (nlftp.mlit.go.jp/) in ArcGIS 10.7.1. For each statistic, forward selections based on the deviance information criterion (DIC) were conducted. The relative migration rate and its inverse, landscape resistance, of each edge were estimated from the selected models. To evaluate the estimates, the correlations between genetic distances and estimated landscape resistance (sum of edges between populations) were calculated by Mantel tests. Future changes in gene flow were inferred under the climate data projected in the representative concentration pathway for high greenhouse gas concentration scenarios (RCP8.5) in IPCC 5th Assessment [31]. Variables based on flow rate and water temperature estimates from 2081 to 2100 were substituted into the BGR model derived from *D*_PS_ to estimate the future gene flow.

**Table 1.** Landscape variables considered in the present study.

| Variable | Description | Hypothesis / Ecological importance | Ranges |
| --- | --- | --- | --- |
| Summer water temperature <sup>A</sup> | Mean water temperature from July to August (July 2019 to August 2019) [°C] | Streams with low summer water temperatures are suitable for *C. nozawae* occupancy/survival [17,20,21] and therefore migration may also occur frequently in these streams. | 8.32–14.2 |
| Water temperature fluctuation <sup>A</sup> | Standard deviation of the water temperature in one year (September 2018 to August 2019) | Thermally stable streams can be suitable for migration. | 1.97–5.09 |
| Drought water discharge <sup>BC</sup> | Flow rate on the day when the flow is 355th highest in one year [m<sup>3</sup>/s] (September 2018 to August 2019) | Drought water discharges, which particularly reflect the environmental heterogeneity created by groundwater [37], ensure opportunities to colonize throughout the year. | 0.07–6.24 |
| Flow fluctuation <sup>AB</sup> | Coefficient of variation in the daily flow rate in one year (September 2018 to August 2019) | Hydrologically stable streams can be suitable for migration. | 0.31–0.65 |
| Edge length | Length of edges [km] | Isolation by distance [38] | 0.05–7.19 |
| Slope | Mean gradient of the edge, i.e., the elevation range divided by edge length | Fish movement and migration are often impeded on steep slopes [39] | 4.15–56.5 |
| Strahler's stream order <sup>C</sup> | Strahler's stream order of the edge | Even in cold-water fish, the mainstem may function as a corridor that facilitates connectivity among populations [9]. | 1–4 |
| Shreve's stream order <sup>BC</sup> | Shreve's stream order (link magnitude), i.e., the numbers of confluence points upstream, at the midpoint of the edge | Given the dendritic arrangement and asymmetry of stream networks, sections with more confluence points upstream may increase the number of migrants passing through [32]. | 1–62 |
| Catchment area <sup>BC</sup> | Cumulative area of the catchment calculated at the midpoint of the edge [km<sup>2</sup>] | Sections with larger catchment areas may have more migrants passing through, based on the same principle as that of the stream order. Or, conversely, streams with larger catchment areas have been identified to have lower *C. nozawae* occupancies [21] and therefore it is also possible that less migration occurs in sections with larger catchment areas. | 0.06–2.96 |
| Direction | Whether the gene flow is toward the upstream (0) or downstream (1) direction | Most stream organisms have higher migration rates in the downstream direction than in the upstream direction [40]. | 0 or 1 |

Variables with the same letters (A, B, C) have high correlations ( $|r| > 0.7$ ); these variables were not included in the same model.

## Results

The average *G*_ST_ was 0.029 (ranged from 0.000–0.047; table S2) and *D*_PS_ was 0.087 (ranged from 0.050–0.135). From the forward selection, Shreve’s stream order, water temperature fluctuation, slope, and edge length were selected for *G*_ST_, and Strahler’s stream order and summer water temperature were selected for *D*_PS_, in this order (tables 2 and S3). In both cases, the first variable added to the model was the stream order, and the second was the water temperature, but of different types between *G*_ST_ and *D*_PS_ (Shreve’s or Strahler’s; summer water temperature or water temperature fluctuation). We could not determine which type of variables precisely affected gene flow because they are highly correlated, but these results show the importance of the stream order and water temperature on the strength of gene flow. The stream orders had a positive effect on gene flow, while the water temperature fluctuation or summer water temperature had a negative effect. The effect (*β*) of the water temperature on gene flow was higher in *D*_PS_ than in *G*_ST_. In *G*_ST_, the slope and edge length were also selected and had negative effects (table 2), indicating their relevance to long-term gene flow. Geographically, the southern upstream area had generally higher landscape resistance (lower gene flow) than the mainstem, while in the northern volcanic area, landscape resistance was not so high even upstream (figure 1). The Mantel tests between the genetic distances and estimated landscape resistance suggested significant relationships, and the correlations were much higher than those between the genetic and geographic distances (r = 0.46, *p* < 0.05 for *G*_ST_; r = 0.60, *p* < 0.01 for *D*_PS_; figure 2). The future prediction indicated that the landscape resistance would increase overall from the current levels. Some sections in the mainstem and in the upper reaches in volcanic areas were estimated to exhibit as high landscape resistance levels as the present southern upstream area. The southern upstream area was projected to display very high resistance.

**Figure 2.**
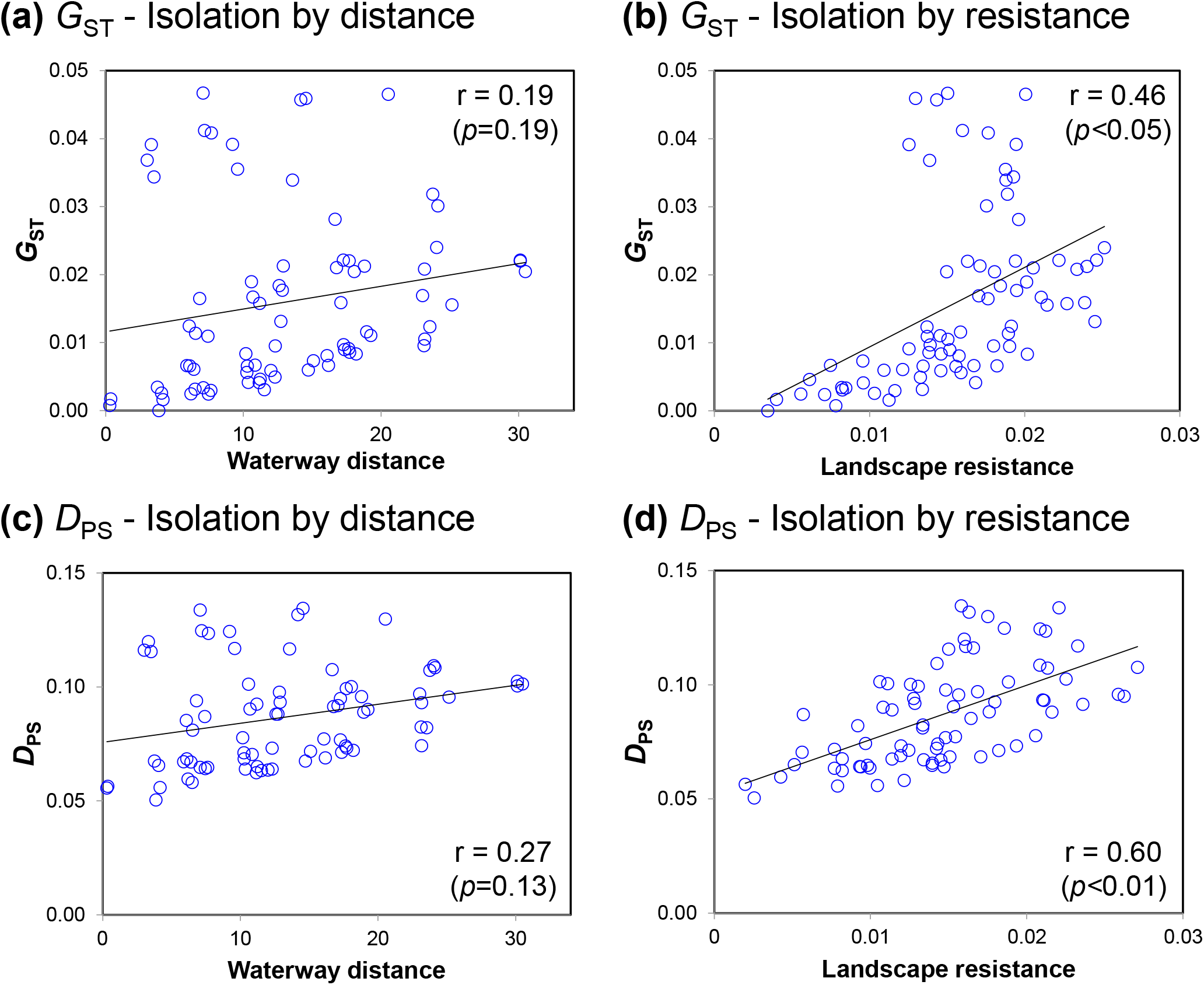
Isolation by distance and isolation by resistance. The relationship between pairwise genetic distance and cumulative landscape resistance between populations (b, d) is compared to the relationship with simple waterway geographic distance (a, c). The case of *G*_ST_ (a, b) and *D*_PS_ (c, d) are shown.

**Table 2.** Selected models explaining the strength of gene flow of the edges. Estimated *β* values (median) and their 95% credible intervals are displayed.

| Variables | $G_{ST}$ | $D_{PS}$ |
| --- | --- | --- |
| (Intercept) | 5.84 [5.65, 5.98] | 6.06 [4.94, 6.85] |
| Shreve's stream order | 4.51 [3.96, 5.19] | -- |
| Strahler's stream order | -- | 2.50 [1.72, 3.15] |
| Water temperature fluctuation | -0.48 [-0.61, -0.35] | -- |
| Summer water temperature | -- | -1.42 [-2.49, -0.38] |
| Slope | -0.94 [-1.15, -0.67] | -- |
| Edge length | -0.42 [-0.54, -0.32] | -- |

## Discussion

In this study, we succeeded in modeling and future predicting of gene flow of *C. nozawae* in the stream network. The modeled landscape resistance well explained the genetic distances (table 2); the strength of gene flow could be largely explained by landscape variables.

It was a somewhat unexpected result that the stream order was identified as the variable with the strongest effect on gene flow. In dendritic stream structures, stream organisms accumulate near confluence points even if random walks are assumed [32], and their passing through downstream edges may result in higher gene flow in higher-order streams. Summer water temperature (or water temperature fluctuation in *G*_ST_) negatively affected gene flow. This is probably because streams with high summer water temperatures and large fluctuations are not suitable environments for *C. nozawae* [21], making successful dispersal difficult. The model from *G*_ST_ also included the slope and edge length, but the *D*_PS_ model did not. We found that topography and distance affected the formation of the long-term population structure as in many other systems, but that most of the current gene flow can be explained by the stream order and water temperature. While the effects of water temperatures have increased from the past to the present, the distribution of landscape resistance has not yet changed extensively (figure 1). The upstream-downstream direction did not affect gene flow, probably because environmental conditions influence the direction of gene flow [17].

Under the RCP8.5 scenario, reduced gene flow and increased landscape resistance across the watershed were predicted (figure 1d). Currently, the studied species exhibits a clear genetic structure only in the southern area (figure S3). The prediction that the northern area will have the same level of gene flow as the present southern area indicates that each tributary within the watershed may experience genetic fragmentation in the future. Nevertheless, gene flow in the northern area was expected to be maintained spatially continuously to some extent, indicating that streams with volcanic watersheds are important for ensuring population connectivity under climate change. A previous study suggested that streams with low summer temperatures behave as source habitats in the watershed [17]. Our study showed that these streams may serve not only as source habitats but also as dispersal pathways in the watershed.

Inoue and Berg [12] considered landscape resistance to be the inverse of the species distribution model (SDM) estimates and predicted that an increased landscape resistance would reduce the gene flow of freshwater bivalves in the future. This is a valuable study that attempts to predict future changes in gene flow; however, it is known that the habitat suitability maps created by SDMs provide very poor estimates of genetic resistance, because of the conceptual differences between habitat selection and entire gene flow [33,34]. Actually, in *C. nozawae*, the SDM created in Suzuki K et al. [21] indicated that the catchment area, analogous to the stream order, had a negative effect on the occurrence of this species, in contrast to the gene flow characteristics estimated in our study. Therefore, genetic population connectivity should be considered separately from habitat suitability.

The present study is novel in that gene flow was modeled using landscape variables identified from genetic data and including water temperature. Our results showed that gene flow in the cold-water sculpin is expected to dramatically decrease in a changing climate. To obtain more robust results, it would be desirable to increase the number of sampling populations. This study has further development potential. For example, demography simulations using inferred landscape resistance [35] could reveal population viability. Also, by combining with habitat quality analyses such as SDMs, population connectivity could be quantified for more detailed predictions from the viewpoint of habitat availability [36]. We hope that riverscape genetics modeling will be applied to predict the consequences of environmental changes on a variety of freshwater organisms.

## Data accessibility

Genetic and environmental data generated in this study were deposited at Figshare [41].

## Author contributions

Conceptualization, S.N. and F.N.; Data curation, S.N. and S.K.H.; Formal analysis, S.N.; Funding acquisition; F.N.; Investigation, S.N., H.S., A.M., and S.K.H.; Methodology, S.N., H.S., M.N., and Y.S.; Project administration, F.N.; Resources, S.N., M.N., A.M., and Y.S.; Software, S.N. and H.S., A.M., and S.K.H.; Supervision, M.N., Y.S. and F.N.; Validation, S.N. and F.N.; Visualization, S.N.; Writing - original draft, S.N. and H.S.; Writing – review & editing, S.N., H.S., M.N., A.M., S.K.H., Y.S., and F.N.

