## Appendix for "Inferring future changes in gene flow under climate change in riverscapes"

Appendix 1. Details of sampling sites. (Figure S1–S2, Table S1)

Appendix 2. Full description of the genetic analysis.

Appendix 3. Full description of the estimation of flow rate and water temperature.

Appendix 4. Supplementary Results. (Figure S3, Table S2–S3)

**Appendix 1.** Details of sampling sites.

**Figure S1** Sampling localities. The blue network indicates the rivers belonging to the Ishikari River system, which has the second largest watershed in Japan and includes the Sorachi River. The present study was conducted upstream of Lake Kanayama, which was formed by the construction of the Kanayama Dam. The node labels correspond to the population IDs listed in Table 1.

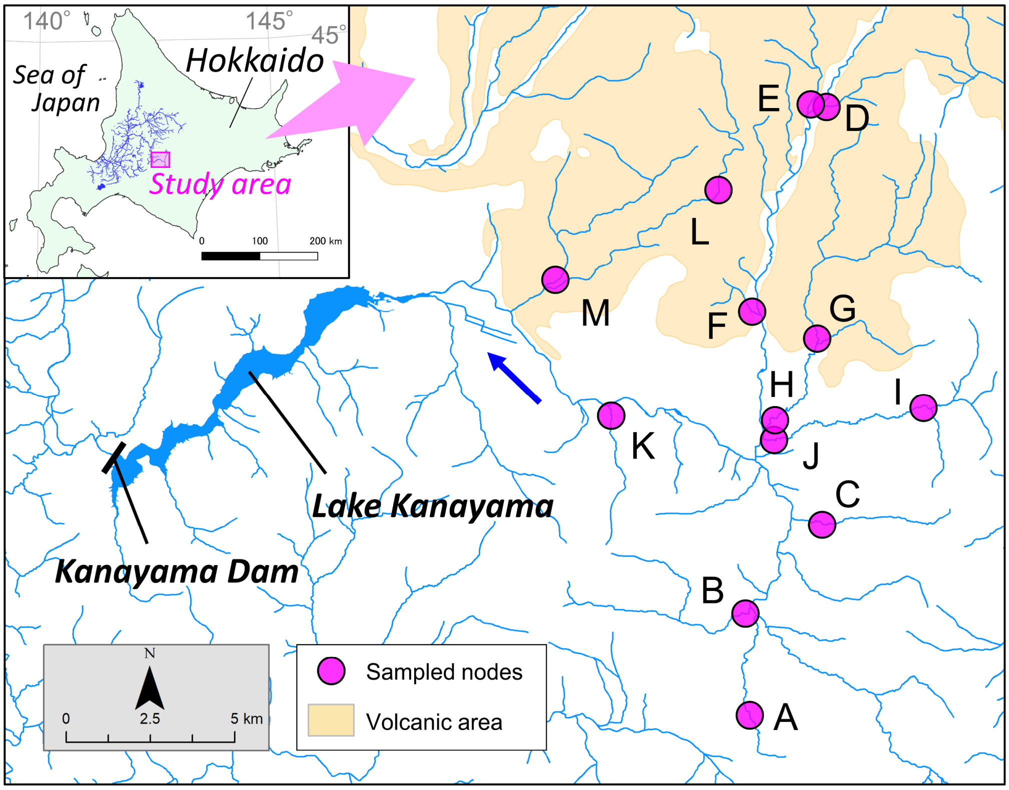

**Table S1** Details of sampling nodes and genetic diversity.

| Pop ID | Name of river | Longitude (E°) | Latitude (N°) | Elevation (m) | N | $H_E$ | $F_{IS}$ |
|---|---|---|---|---|---|---|---|
| A | Tomamu R. | 142.670 | 43.057 | 526 | 32 | 0.258 | 0.013 |
| B | Kinnosawa R | 142.672 | 43.085 | 466 | 22 | 0.265 | -0.002 |
| C | Shikerebenaizawa R | 142.699 | 43.107 | 451 | 31 | 0.255 | 0.014 |
| D | Ozawa R. | 142.703 | 43.218 | 511 | 30 | 0.267 | -0.006 |
| E | Ehoroakanbetsu R. | 142.699 | 43.218 | 510 | 34 | 0.271 | -0.004 |
| F | Koya-no-sawa R. | 142.678 | 43.163 | 445 | 21 | 0.262 | -0.003 |
| G | Pankeyara R. | 142.699 | 43.155 | 450 | 32 | 0.265 | -0.004 |
| H | Pankeyara R | 142.680 | 43.133 | 414 | 32 | 0.272 | 0.004 |
| I | Peiyurushiebe R. | 142.739 | 43.136 | 498 | 32 | 0.265 | 0.011 |
| J | Peiyurushiebe R. | 142.680 | 43.129 | 413 | 32 | 0.265 | 0.023 |
| K | Ecchudantai-no-sawa R. | 142.624 | 43.137 | 387 | 32 | 0.264 | -0.012 |
| L | Ikutora R. | 142.665 | 43.196 | 509 | 14 | 0.241 | -0.002 |
| M | Kuma-no-sawa R. | 142.603 | 43.173 | 376 | 32 | 0.265 | 0.007 |

Pop ID, population ID of sampling nodes; N, number of sampled individuals; $H_E$, expected heterozygosity; $F_{IS}$, fixation index. $H_E$ and $F_{IS}$ were calculated in the *populations* command in STACKS.

**Figure S2** Spatial distribution of landscape variables. The edges are colored by the values of each variable.

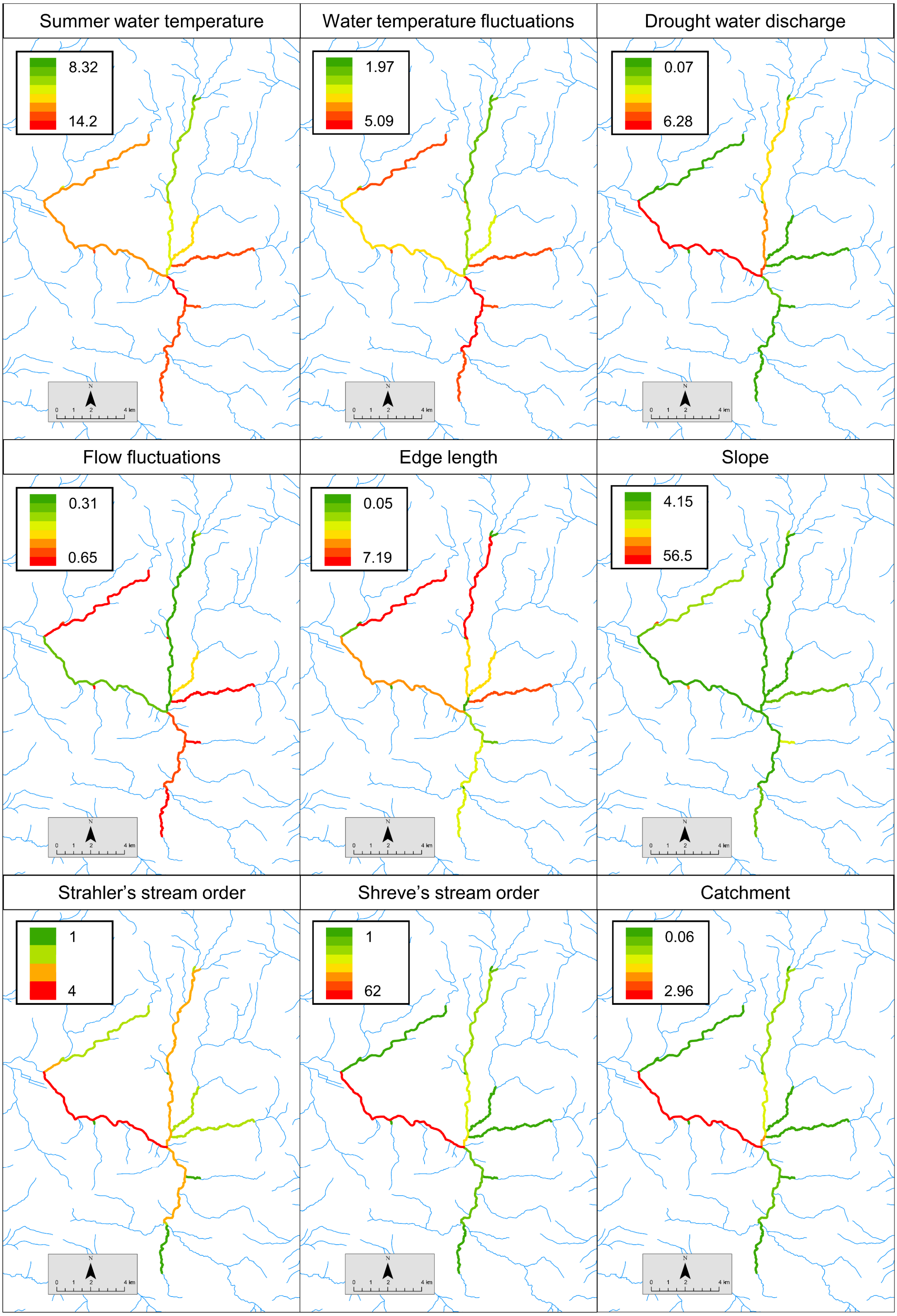

**Appendix 2.** Full description of the genetic analysis.

Total DNA was extracted using the QIAGEN DNeasy Blood and Tissue Kit (QIAGEN Inc.). To obtain the genetic data, we used multiplexed ISSR genotyping by sequencing (MIG-seq; Suyama and Matsuki 2015), a technique in which loci between two microsatellite regions are amplified by PCR and next-generation sequencing. A MIG-seq library preparation and read quality filtering were performed following the protocol described in Suyama et al. (2022), with the modification that two runs were conducted and the obtained data were combined after quality filtering. In addition, quality filtering was performed on 71 bases with six 5'-end bases and three 3'-ends bases removed. After the quality filtering, SNP selection was performed using STACKS 2.41 (Catchen et al. 2013). First, the reads were grouped to each locus using the *ustacks*, *cstacks*, *sstacks*, *tsu2bam*, and *gstacks* commands with the following parameters recommended by Paris et al. (2017): minimum depth option creating a stack ($m$) = 3, maximum distance between stacks ($M$) = 2, maximum mismatches between loci when building the catalog ($n$) = 2, and number of mismatches allowed to align secondary reads ($N$) = 4. From the derived dataset of assembled loci, SNPs were detected using the *populations* commands under the following criteria: only loci present at a rate of more than 80% of individuals within all populations were extracted (-p 13 -r 0.8); the minimum minor allele frequency was 5% (--min-maf 0.05); sites showing excess heterozygosity were removed (--max-obs-het 0.6); and the output was limited to one SNP per locus (--write-single-snp). After filtering, 212 SNPs were obtained.

Genetic differentiation among populations was assessed by $G_{ST}$ (Nei 1973) and $D_{PS}$ (Bowcock et al. 1994). $D_{PS}$ is the genetic distance based on the dissimilarities of population allele pools and reflects gene flow over a shorter timescale (approximately 10 generations; Landguth et al. 2010; Leroy et al. 2017), whereas $G_{ST}$ is assumed to reflect long-term gene flow (Holsinger & Weir 2009). $G_{ST}$ was calculated using GenAlEx 6.51 (Peakall and Smouse 2012), and $D_{PS}$ was calculated using the package graph4lg (Savary et al. 2021) in R 3.6.0 (R Core Team 2019).

Gene flow of *C. nozawae* in the stream network was estimated by using the novel bidirectional geneflow in riverscapes (BGR) model (White et al. 2020). In this model, bidirectional gene flow in a stream network can be modeled using genetic distance matrices as a response and landscape variables as explanatory variables, rigorously accounting for the spatial autocorrelation structure of dendritic stream networks. Specifically, the nearly homogeneous stream segments (delimited by nodes that are sampling sites or major tributary confluences) were defined as edges, and the relative migration rate (edge passability; $w_{ij}$) of each edge linking nodes $i$ and $j$ was estimated as a function of $k$ landscape variables ($x_{ij1}$, $x_{ij2}$, …, $x_{ijk}$) and the corresponding parameters ($\beta_1$, $\beta_2$, …, $\beta_k$) as:

$$w_{ij} = \exp\,(\beta_0 + \sum_k \beta_k x_{ijk})$$

where $\beta_0$ is the intercept term. Here, all landscape variables were standardized from 0 to 1. The posterior distribution of parameters $\beta_0$, $\beta_1$, …, $\beta_k$ was estimated by a Markov Chain

Monte Carlo (MCMC) sampler.

We used $G_{ST}$ and $D_{PS}$ as genetic distances and 10 landscape variables described in Table 1 as covariates $x_{ijk}$. All variables except direction are symmetric. Edge length indicates the length of the edge and corresponds to the isolation by distance (IBD) hypothesis, but note that the model assumes IBD constantly as the number of passing edges.

For each summary statistic, forward selections were conducted based on the deviance information criterion (DIC). Variables were added until the DIC no longer decreased by 7 or more (Cain & Zhang 2018). At each step of the forward selection, the variables that were highly correlated (Pearson's $r > 0.7$) with other variables already included in the model were not added to the model. Models with fewer than four variables were run for 50,000 MCMC iterations and parameters were estimated after 25,000 burn-in. Models with four or more variables were run for 100,000 iterations including 50,000 burn-in. After the final model was identified, we conducted a long run with 500,000 iterations including 200,000 burn-in, to accurately estimate the $\beta$ values and 95% credible intervals. Landscape resistance, calculated as the inverse of $w_{ij}$ of each edge, was estimated and mapped from the selected models. To evaluate the estimates, the correlations between genetic distances and estimated landscape resistance (sum of edges between populations) were calculated by Mantel tests with 9999 permutations, and compared to the correlations between genetic distances and waterway geographical distance. Future landscape resistance was inferred by substituting future water temperature (mean of predicted variables from 2081 to 2100) into the final BGR model derived from $D_{PS}$.

To discuss the results with genetic structure, we also conducted a conventional STRUCTURE analysis (Pritchard et al. 2000). The STRUCTURE settings were the admixture and allele frequency correlated model with previous sampling location information (LOCPRIOR; Hubisz et al. 2009). The algorithm was run 10 times for each K from 1 to 10 with a burn-in of 20,000 followed by 30,000 MCMC replicates. The program CLUMPAK (Kopelman et al. 2015) was then used to summarize the results for each K. STRUCTURE HARVESTER (Earl and vonHoldt 2012) was employed to calculate the probability of the data for each K (LnP(D); Pritchard et al. 2000), the corresponding standard deviation, and the ΔK (Evanno et al. 2005).

**Appendix 3.** Full description of the estimation of flow rate and water temperature.

The flow rate and water temperature were estimated according to Suzuki H et al. (2021) and Nakatsugawa and Suzuki (2022). In the following, a summary of the methodology is described according to these studies.

From March 2018 to September 2019, the water level and temperature were measured in two subwatersheds where Quaternary Pleistocene pyroclastic flow deposits are distributed (sites E and M in Figure S1) and in two subwatersheds where Mesozoic rocks are distributed (sites B and J in Figure S1). Manning's roughness coefficients, determined from field surveys, were applied to Manning's mean velocity formula to convert the daily mean water levels to flow rates. These flow rates and water temperatures were used to verify the reproducibility of the values calculated by the model described below. The reproducibility was also verified using publicly available data provided by the Ministry of Land, Infrastructure, Transport and Tourism of Japan (Water Information System; www1.river.go.jp/) on the inflow into Lake Kanayama and the water temperatures in the lower reaches of the Lake Kanayama watershed.

Daily precipitation, snowmelt, and evapotranspiration were calculated on each 1 km mesh using an atmospheric land surface process model (Usutani et al. 2005), in which snow accumulation and snowmelt processes are incorporated. As the input meteorological data, downscaled data from the Hokkaido Branch of the Japan Weather Association was used to reproduce the current conditions (March 2018 to September 2019), and downscaled data from Ueda et al. (2020) was used for future climate simulations (2081 to 2100).

The sum of the calculated precipitation and snowmelt minus evapotranspiration was used as the effective precipitation, and the slope runoff was obtained by summing the runoff at each 1 km mesh using the four-layered tank model (Sugawara 1979). In areas where pyroclastic flow deposits are distributed (hereafter, "volcanic areas"), a bottom outlet was added to the fourth tank of the tank model, and water infiltrating through this added bottom outlet was assumed to flow out at the mainstem of the Sorachi River, to account for the subsurface flow of groundwater. The tank model parameters were determined for each subwatershed using the SCE-UA method (Duan et al. 1992), one of the common optimization methods, and were adjusted through trial and error. The geology of the Lake Kanayama watershed was classified into volcanic areas and other areas (in which Mesozoic rocks were distributed), and the tank model parameters for each subwatershed were given corresponding to each geology type to reflect the runoff characteristics according to the geological conditions. The slope runoff was summed, and total discharge was calculated by streamflow routing using the kinematic wave equation.

Different water temperatures were given for each outflow component from each tank in the tank model. For the first tank, the water temperature was set to 273.15 K during snow accumulation and set equal to the daily average temperature (with a lower limit of 273.15 K) during non-snow accumulation. For the fourth tank, the annual mean temperature plus 1.5 (K) was set as the groundwater temperature in reproducing the current conditions; this value was also assumed to remain in future climate simulations. The water temperatures in the second tank and the third tank were calculated by proportionally dividing the water temperatures of the outflows from the first tank and the fourth tank. Different proration coefficients were given for the second tank and the third tank, and for the volcanic areas and other areas, to allow the observed water temperatures to be well reproduced. The water temperature fluxes of these slope runoff and the air-surface water temperature fluxes derived from the heat balance equation for the water surface were summed, and the water temperature fluxes were then tracked along the streamflow using the kinematic wave equation. Here, water temperature flux represents the product of flow rate and water temperature, in units of $m^3 s^{-1} K$. The sum of the water temperature fluxes was divided by the flow rate to estimate the stream water temperature at each mesh.

Under the above conditions, the reproduction of flow rate and water temperature under the current conditions was verified and confirmed to be well reproduced. Water temperatures for future climate simulations were then calculated.

**Appendix 4.** Supplementary Results.

**Table S2** Pairwise $G_{ST}$ (below diagonal) and $D_{PS}$ (upper diagonal) matrices among the sampling nodes. Row and column headings (A–M) correspond to population IDs in Table and Figure S1.

| | A | B | C | D | E | F | G | H | I | J | K | L | M |
|---|---|---|---|---|---|---|---|---|---|---|---|---|---|
| A | | 0.085 | 0.117 | 0.109 | 0.107 | 0.108 | 0.091 | 0.093 | 0.096 | 0.088 | 0.095 | 0.102 | 0.093 |
| B | 0.012 | | 0.116 | 0.100 | 0.099 | 0.101 | 0.090 | 0.094 | 0.088 | 0.081 | 0.093 | 0.109 | 0.092 |
| C | 0.035 | 0.034 | | 0.135 | 0.132 | 0.134 | 0.125 | 0.120 | 0.124 | 0.116 | 0.124 | 0.130 | 0.117 |
| D | 0.030 | 0.020 | 0.046 | | 0.056 | 0.064 | 0.072 | 0.065 | 0.073 | 0.063 | 0.074 | 0.101 | 0.082 |
| E | 0.032 | 0.022 | 0.046 | 0.002 | | 0.065 | 0.068 | 0.070 | 0.071 | 0.062 | 0.077 | 0.101 | 0.074 |
| F | 0.028 | 0.019 | 0.047 | 0.002 | 0.003 | | 0.065 | 0.068 | 0.071 | 0.066 | 0.078 | 0.097 | 0.077 |
| G | 0.021 | 0.017 | 0.041 | 0.007 | 0.006 | 0.003 | | 0.050 | 0.064 | 0.056 | 0.068 | 0.082 | 0.069 |
| H | 0.021 | 0.016 | 0.039 | 0.005 | 0.007 | 0.003 | 0.000 | | 0.058 | 0.056 | 0.067 | 0.090 | 0.064 |
| I | 0.021 | 0.013 | 0.039 | 0.009 | 0.009 | 0.006 | 0.004 | 0.003 | | 0.060 | 0.073 | 0.096 | 0.072 |
| J | 0.018 | 0.011 | 0.037 | 0.003 | 0.004 | 0.003 | 0.002 | 0.001 | 0.002 | | 0.068 | 0.089 | 0.064 |
| K | 0.022 | 0.016 | 0.041 | 0.009 | 0.010 | 0.008 | 0.007 | 0.006 | 0.009 | 0.007 | | 0.098 | 0.067 |
| L | 0.022 | 0.024 | 0.047 | 0.020 | 0.022 | 0.017 | 0.010 | 0.011 | 0.016 | 0.012 | 0.018 | | 0.087 |
| M | 0.021 | 0.016 | 0.034 | 0.012 | 0.011 | 0.008 | 0.007 | 0.005 | 0.008 | 0.006 | 0.007 | 0.011 | |

**Table S3** Models considered in the forward selection process. All models include an intercept term, although omitted in the table. The boldface indicates the model displaying the lowest DIC at each step, while the italics indicate the model displaying the lowest DIC but not meeting the criteria to include the variable (i.e., DIC decrease by 7 or more). WTS = summer water temperature, WTF = water temperature fluctuation, FlowD = drought water discharge, FlowF = flow fluctuations, Length = edge length, Strahler = Strahler's stream order, Shreve = Shreve's stream order, Catchment = catchment area.

| $D_{PS}$ | | $G_{ST}$ | |
|---|---|---|---|
| Model | DIC | Model | DIC |
| (one variable) | | (one variable) | |
| WTS | -15073 | WTS | -20045 |
| WTF | -15073 | WTF | -20159 |
| FlowD | -15069 | FlowD | -20409 |
| FlowF | -15070 | FlowF | -20487 |
| Length | -15054 | Length | -19740 |
| Slope | -15057 | Slope | -20087 |
| Strahler | **-15080** | Strahler | -20429 |
| Shreve | -15070 | Shreve | **-20667** |
| Catchment | -15070 | Catchment | -20626 |
| Direction | -15065 | Direction | -19859 |
| (two variables) | | (two variables) | |
| Strahler+WTS | **-15089** | Shreve+WTS | -20772 |
| Strahler+WTF | -15086 | Shreve+WTF | **-20787** |
| Strahler+FlowF | -15086 | Shreve+Length | -20713 |
| Strahler+Length | -15085 | Shreve+Slope | -20743 |
| Strahler+Slope | -15088 | Shreve+Direction | -20753 |
| Strahler+Direction | -15080 | | |
| (three variables) | | (three variables) | |
| Strahler+WTS+Length | -15090 | Shreve+WTF+Length | -20812 |
| Strahler+WTS+Slope | *-15091* | Shreve+WTF+Slope | **-20860** |
| Strahler+WTS+Direction | -15086 | Shreve+WTF+Direction | -20824 |
| | | (four variables) | |
| | | Shreve+WTF+Slope+Length | **-20927** |
| | | Shreve+WTF+Slope+Direction | -20050 |
| | | (five variables) | |
| | | Shreve+WTF+Slope+Length+Direction | *-20931* |

**Figure S3** Population structure inferred in STRUCTURE. (a) Barplots displaying the proportion of the membership coefficient in the inferred subpopulations (clusters) at K = 2, 3, 4, and 6 for all individuals. The letters at the bottom indicate the population IDs. (b) The mean values of the posterior probability of the data (LnP(D)) from 10 runs for each K. (c) The values of ΔK at each K.

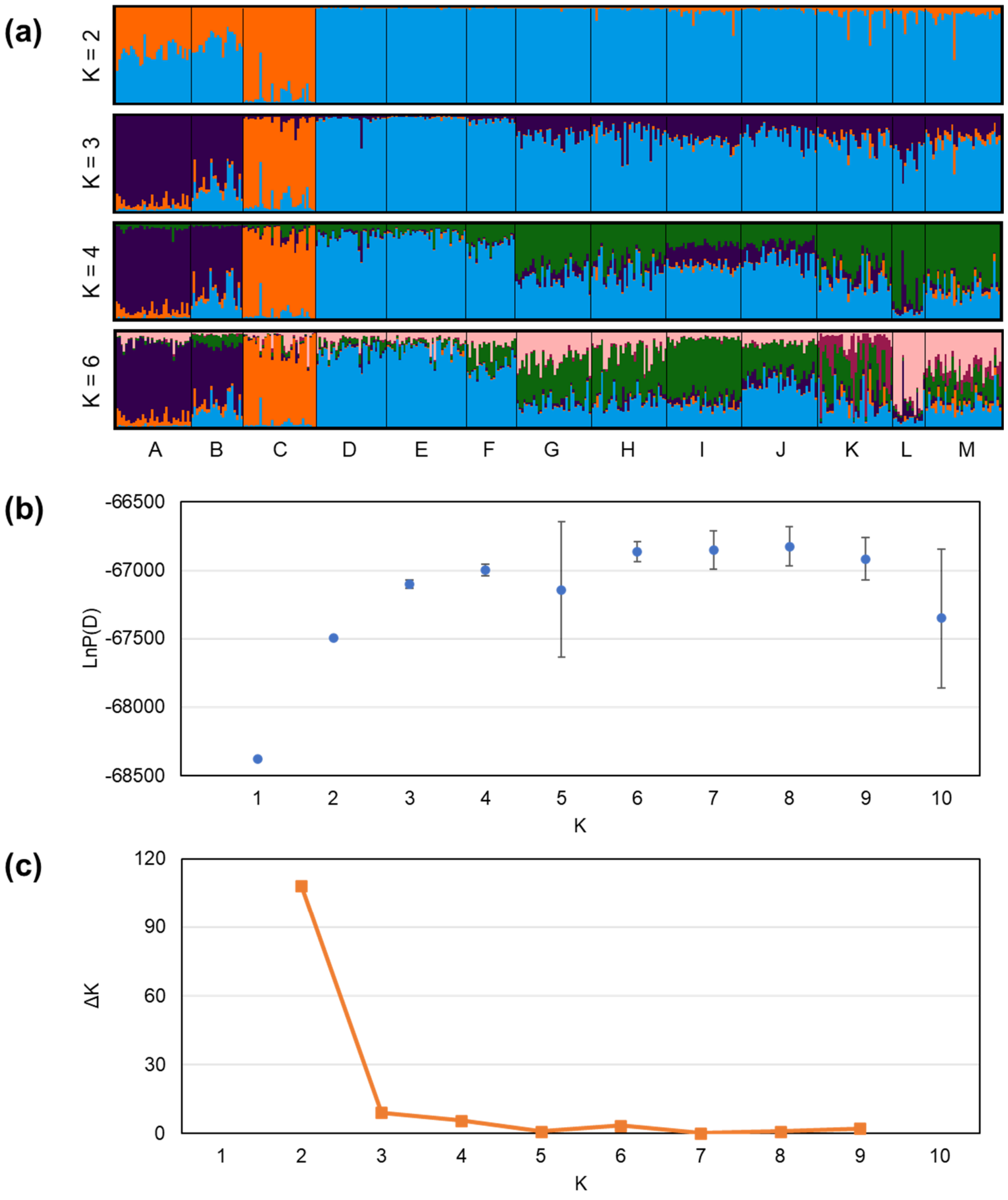
